# Genome-wide analysis of the apple family 1 glycosyltransferases identified a flavonoid-modifying UGT, MdUGT83L3, which is targeted by MdMYB88 and contributes to stress adaptation

**DOI:** 10.1101/2021.02.01.429094

**Authors:** Yanjie Li, Pan Li, Lei Zhang, Jing Shu, Michael H Court, Zhuojing Sun, Lepu Jiang, Chengchao Zheng, Huairui Shu, Bingkai Hou, Lusha Ji, Shizhong Zhang

## Abstract

Nowadays, the plant family 1 UDP-glycosyltransferases (UGTs) are more and more investigated owing to their contribution to the plant secondary metabolism and diverse biological roles. Apple (Malus domestica) is one of the most widely cultivated fruit trees with great economical importance. However, poor knowledge is known about the apple UGTs. In this study, we identified 229 members of family 1 through a genome-wide analysis of the apple UGTs, which were clustered into 18 groups, from A to R. We also performed detailed analysis to 34 apple UGTs with quantitavive RT-PCR, and discovered a number of stress-regulated UGTs. Among them, we characterized the role of MD09G1064900, also named as MdUGT83L3, which is significantly induced by salt and cold. In vivo analysis showed that it has high activity towards cyanidin, and moderate activity towards quercetin and keampferol. The transgenic callus and regenerated apple plants overexpressing MdUGT83L3 showed enhanced tolerance to salt and cold treatments. Overexpression of MdUGT83L3 also increased anthocyanin accumulation in the callus tissues and enhanced ROS clearing upon exposed to salt and cold stresses. Futhermore, via yeast-two-hybrid assay, EMSA and CHIP analysis, we also found that MdUGT83L3 could be directly regualted by MdMYB88. Our study indicated that MdUGT83L3, under the regulation of MdMYB88, plays important roles in salt and cold stress adaptation via modulating flavonoid metabolism in apple.

## Introduction

Plants produce numerous and diverse structurally specialized metabolites—which is also called “natural products” and fulfill crucial ecological functions. Apple (*Malus domestica*) is one of the most widely cultivated fruit trees in temperate regions with great economical importance for landscaping and reforestation. As a kind of popular and delicious fruit, apple contains richful natural products, especially flavonoids, such as flavonoids, dihydrochalcones and dihydrochalcones ^1,2^, which contributes to the appeal colors and the nutrition of the friuts. In traditional Chinese medicine, *Malus hupehensis* had been used to treat inflammatory diseases due to its rice flavonoids, phenols, and amino acids ^3^. It is also reported that apple can reduce cancer risk with flavonoids ^4^.

Flavonoids are a large class of phenolic compounds synthesized in plants and are closely related to normal plant development and adaptation to the environment. These compounds, which includes flavonols, flavones, isoflavones, and anthocyanins, are mainly synthesized through the phenylpropane metabolic pathway. First, chalcone synthase (CHS) catalyzes 4-coumarin-CoA and malonyl CoA into chalcone, which then goes through a series of enzymatic reactions and becomes dihydroflavonols (including dihydrokaempferol and dihydroquercetin). The dihydroflavonols, on one side, are catalyzed into quercetin and kaempferol under the action of flavonol synthase (FLS), and on the other branch are transferred into anthocyanins by hydroflavonoid reductase (DFR) ^5^. Finally, the quercetin, kaempferol, and anthocyanins are conjucted with various sugar groups, which become more stable and soluble, thus facilitate their transportation and storage in plant cells in the form of glycosides ^6,7^.

Flavonoids exist mostly as their glycosides in plants, which are catalyzed by family 1 glycosyltransferases, also designated as UDP-glycosyltransferases (UGTs) ^8,9^. Although many flavonoid-modifying UGTs have been identified, they tend to be under different regulations or have specific roles. For instance, the Arabidopsis subfamily UGT79B1, 79B2 and 79B3 were found to modify anthocyanins, however, UGT79B1 only recognizes 3-O-glucosylated anthocyanidins/flavonols, while UGT79B2 and 79B3 prefer anthocyanidins rather than 3-O-glucosylated anthocyanidins, which were directely controled by CBF1 ^10,11^. Another small homologous group UGT78D1, 78D2 and 78D3 also modify flavonoids. Of these, UGT78D2 catalyzes the transfer of UDP-glucose onto cyanidin, kaemferol and quercetin ^12^, While UGT78D1 and 78D3 are specific for flavonol aglycones only ^8^. The maize UDP glucose:flavonoid glucosyltransferase (UFGT) BRONZE is the first identifed UGT gene in plants. It is specifically expressed in the endosperm and responsible for seed pigmentation via catalyzing the 3-O-glucosylation of flavonols or anthocyanidins ^13^. The second characterized maize UFGT gene UFGT2, shows high activity towards kaemferol and quercetin, and contributes to abiotic stress tolerance ^14^.

The R2R3-MYB TF family, which is the largest MYB subfamily, has been demonstrated to act as the main regulator involved in flavonoid biosynthesis in many plant species. Previous studies have thoroughly elucidated the anthocyanin biosynthesis pathway that is regulated by the conserved MBW ternary complex, including the R2R3-MYB, basic helix-loop-helix (bHLH), and WD40-repeat subunits in higher plants. The bHLH proteins bind to MYB and WD40 to form the MBW complex, which activates the expression of anthocyanin-specific genes, including DIHYDROFLAVONOL 4-REDUCTASAE (DFR), ANTHOCYANIDIN SYNTHASE (ANS), UDP-GLUCOSE: FLAVONOID 3-O-GLUCOSYL TRANSFERASE (UF3GT), BANYULS/anthocyanidin reductase (BAN), TRANSPARENT TESTA19 (TT19), TT12 and AHA10^15^. Till now, many MBW subunits have been identified in higher plants, such as the Arabidopsis MYB factors PRODUCTION OF ANTHOCYANIN PIGMENTATION 1 (PAP1)/MYB75, PAP2/MYB90, MYB113 and MYB114^16,17^, bHLH transcription factors TRANSPARENT TESTA 8 (TT8), GLABRA 3 (GL3), and ENHANCER OF GLABRA 3 (EGL3). However, till now, only one WD40-repeat protein, TRANSPARENT TESTA GLABRA 1 (TTG1), has been found in Arabidopsis ^18,19^.

The regulation of anthocyanin biosynthesis has always been a hot topic in apple. The red colour of apple skin is composed almost exclusively of cyanidin-3-O-galactoside, which is an anthocyanin derivative. A number of studies have revealed the role of MYB factors in the regulation of anthocyanin biosynthesis in apple, especially in recent years. Earlier studies reported that MdMYB1 and its allelic genes act as the central players in regulating anthocyanin biosynthesis and fruit coloration ^20-22^. MYB10 and MdbHLH3, together with BBX1, were reported to regulate the fruit colour by activating the anthocyanin biosynthetic gene DFR^23^. An et al. (2015)^24^ reported that overexpression of MdMYB9 or MdMYB11 promoted both anthocyanin and PA accumulation in apple callus, which was enhanced by MeJA. Well, it is not a unique case. The MdMYB24-like (MdMYB24L), which is induced by JA, was also found to regulate the anthocyanin biosynthesis via JA signaling pathways ^25^. Lately, An et al. (2019)^26^ reported that the MdMYB308L interacted with MdbHLH33 and enhanced its binding to the promoters of MdCBF2 and MdDFR, suggest that MdMYB308L acts as a positive regulator in cold tolerance and anthocyanin accumulation.

The MdMYB88 is a well-studied transcription factor in recent years which has been identified to play pleiotropic roles via controlling diverse target genes in apple. According to former study^27^, MdMYB88 and its paralog MdMYB124 regulate cold tolerance by targeting *COLD SHOCK DOMAIN PROTEIN 3 (MdCSP3)* and *CIRCADIAN CLOCK ASSOCIATED 1 (MdCCA1)* genes, and they also promote anthocyanin accumulation and H_2_O_2_ detoxifification in response to cold stress. MdMYB88 and MdMYB124 were also reported to regulate root xylem development by directly binding MdVND6 and MdMYB46, which thus enhanced drought tolerance ^28^. Lately, the same group further found that MdMYB88 and MdMYB124 could directly target MdCM2, a critical enzyme involved in phenylalanine biosynthesis, and thus significantly affects flavoniod metabolites in apple roots ^28^. Moreover, MdMYB88 and MdMYB124 were also found to control the expression of some other genes such as TIME FOR COFFEE (TIC), which contributes to freezing tolerance by promoting unsaturation of fatty acids in apple ^29^, and MdNCED, a key enzyme involved in ABA biosynthesis ^30^, as well as three BR signaling pathway genes DE ETIOLATED 2 (MdDET2), DWARF 4 (MdDWF4), and BRASSINOSTEROID 6 OXIDASE 2 (MdBR6OX2) (Liu et al., 2021)^31^.

Till now, more than 200 UGTs have been reported in apple^32^. However, only a very limited number of UGTs have been functionally characterized till now. For example, Elejalde-Palmett et al. (2019)^33^ found that members of UGT88F subfamily catalyzes a wide range of flavonoids including phloridzin flavonols, flavones, flavanones, chalcones and dihydrochalcones in vitro. A further research found that MdUGT88F1 mediate the biosynthesis of phloridzin and participates in the plant development and pathogen resistance in apple trees^34^. The MdUGT75B1 and MdUGT71B1 were identified as key UGTs involved in flavonol galactoside/glucoside biosynthesis in apple fruit^35^. Apart from these, most of the apple UGTs remain to be elucidated in both the biochemical and biological levels.

In this study, we characterized a novel and stress-responsive apple UDP-glycosyltransferases MD09G1064900, which has high activity towards anthocynins, and moderate activity towards quercetin and keampeferol. In alignment with the known plant UGTs by blast, it has the highest sequence identity with UGT83L3, and was thus named MdUGT83L3. We also found that MdUGT83L3, which is directly controlled by MdMYB88, is involved in the regulation of flavonoid biosynthesis in response to salt and cold stress.

## Results

### Identification and phylogenetic analysis of the apple family 1 glycosylatransferase

The comprehensive sequencing of the apple genome has greatly facilitated the identification of apple gene families. To identify apple UGT genes, the 44-amino acid sequence that constitutes the conserved PSPG box was used for searching apple UGTs with BlastP against the apple genome database on both NCBI (https://www.ncbi.nlm.nih.gov/) and GDR (https://www.rosaceae.org/species/malus/all). After careful analysis and confirming all the amino sequences contain the PSPG signature motif, we then removed redundant gene IDs encoding the same proteins, and redundant sequences representing spliced versions, we finally got 229 apple UGT genes (Table 1), which is a little less than but in proximity to the UGT numbers (241) reported in previous reports^32^. Phylogenetic analysis is very crucial for effectively analyzing UGT gene functions, and UGTs with relevant roles tend to locate closely (Wilson and Li, 2019). To group the 229 apple UGTs, an unrooted tree was constructed from alignments of the full-length amino acid sequences of the apple UGTs together with 20 Arabidopsis UGTs, including AtUGT71B1, AtUGT72C1, AtUGT73C1, AtUGT74C1, AtUGT75C1, AtUGT76B1, AtUGT78D3, AtUGT79B1, AtUGT82A1, AtUGT83A1, AtUGT84A1, AtUGT85A1, AtUGT86A1, AtUGT87A1, AtUGT88A1, AtUGT89B1, AtUGT90A1, AtUGT91A1, AtUGT91B1 and AtUGT92A; three maize UGTs including GRMZM2G168474_P02, GRMZM5G834303_P01, GRMZM2G162783_P01; one UGT PgUGT95B2 from *Punica granatum L*, one UGT VvUGT85A24 from *Vitis vinifera*, one cassava (*Manihot esculenta*) UGT MeUGT85K4, one *Medicago truncatula* UGT MtUGT85H2, and one UGT in *Catharanthus roseus* CrUGT709C2. The 229 apple UGTs were finally clustered into 18 groups, from A to R (Figure 1A), which is an update to the data in previous study^32^. According to previous studies, the plant UGTs have been put into 18 groups (A to R) as well as two outgroups OG80 and OG81. In our tree, 14 groups (A-N) were clustered with the UGTs from Arabidopsis, *Vitis vinifera, Manihot esculenta, Medicago truncatula* as well as the known apple UGTs (MdUGT71B1 and MdUGT88F4). We also employed group O member GRMZM2G168474_P02, group P members GRMZM5G834303_P01 and CrUGT709C2, group Q member PgUGT95B2, and group R member GRMZM2G162783_P01 for constructing the tree, and got two group O members, five group P members, one group Q member and four group R members. The outgroup members are identified to modify lipids and sterols and their PSPG-box is less conserved. However, we did not found any apple UGTs belonging to the outgroup.

**Figure 1.**
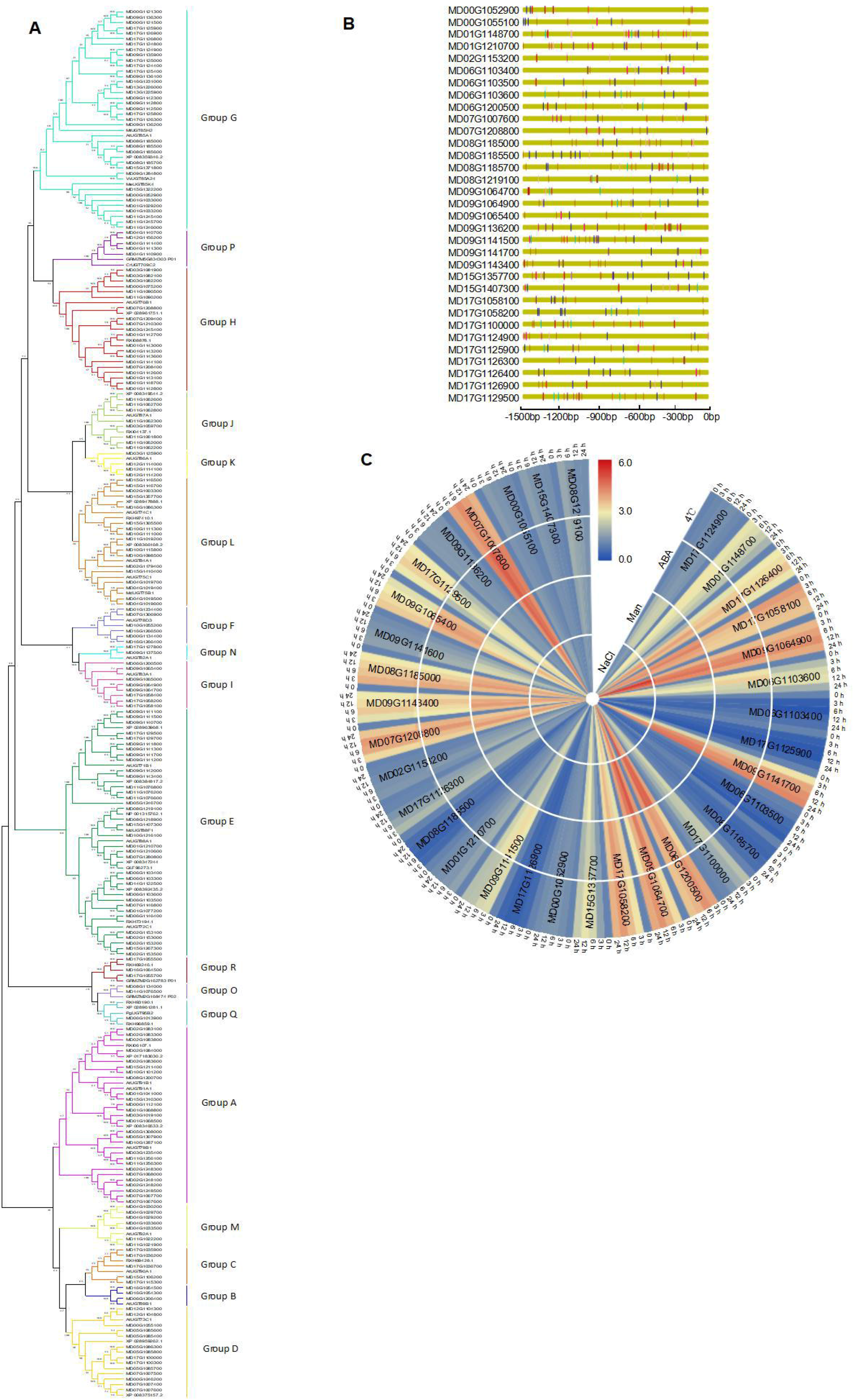
Genome-wide analysis of Family-1 UGTs from apple. **A**. The neighbor-joining tree was created using the MEGA-X program (bootstrap value set at 1,000) with full-length sequences of 229 apple UGTs, 20 Arabidopsis UGTs, three maize UGTs, one UGT (PgUGT95B2) from Punica granatum L, one UGT (VvUGT85A24) from grape, one UGT (MeUGT85K4) from cassava, one UGT (MtUGT85H2) from Medicago, and one UGT (CrUGT709C2) in Catharanthus roseus. Bootstrap values over 50 % are indicated above the nodes. **B**. Analysis of the stress-related cis-element present in the upstream region of 34 apple UGTs. **C**. The 2-week-old apple seedlings were subjected to NaCl, ABA, Mannitol and Cold for 3, 6, 9, 12 hours, respectively, and the expression of these 34 UGTs were analyzed with quantitative real-time PCR.

### Analysis of a subset of apple family 1 glycosyltransferase identifies a stress responsive UGT

It is reported in previous studies that plants employ a lot of UGT members to cope with various environmental stresses, which play important roles in stress adaptation ^11,36,37^. Since for apple there is no transcriptome data deposited on the public platform (for example, genevestigator), we thus analyzed the upstream sequences of a number of apple UGTs for stress-regulated elements (Figure 1B). Based on the presence of common stress-related elements, we selected 34 apple UGTs for further analysis. The apple seedlings were exposed to time-course treatment under NaCl, mannitol, cold (4°C) and ABA, respectively, for 0, 3, 6, 12, 24 hours. The expression of all the 34 UGTs were assessed in detail with qRT-PCR (Figure 1C). Among the 34 UGTs, a large number of them are not responsive to these stresses as the blue colour indicated, including MD06G1103500, MD17G1129500, MD08G1185700 and MD08G1185500 *etc*. And a lot of them are moderately induced by these stresses as the indicated in yellow, such as MD06G1103600, MD09G1141500, MD15G1357700, MD17G1129500 *etc*. What interested us is that several genes are significantly upregulated upon these stimuli as indicated in red, such as MD09G1064900, MD17G1058100, MD09G1141700, MD07G1007600 etc, which might be involved in stress response. To make further investigation, here, we performed protein expression and substrate screening for these four genes, and finally, we paid our attention to MD09G1064900 which belongs to UGT83 subfamily. Roles of UGT83 family members have not been reported yet, thus, it would be interesting make some investigation. Further qRT-PCR confirmed that MD09G1064900 is responsive to NaCl, Mannitol, ABA and Cold, and the UGT is significantly induced by NaCl (up to 25 folds), and moderately induced by ABA, mannitol and cold (below 10 folds) (Figure S1A). We also investigated its spatio-temperol expression and observed that the gene could be expressed in various tissues, with relatively abundant expression in stem and lower expression in root and flower (Figure S1B). Furthermore, according to the UGT Nomenclature commitee, MD09G1064900 is named as MdUGT83L3.

### MdUGT83L3 catalyzes the glycosylation of cyanidin, quercetin and kaempferol

To determine the substrate(s) of MdUGT83L3,the recombinant GST-tagged MdUGT83L3 protein were purified, and subjected to the substrate screening towards a broad scope of metabolites in the phenylpropanoid metabolic pathway, including cinnamic acid, ferulic acid, caffeic acid, etc., from the early steps of the phenylpropanoid pathway, and cyanidin, quercetin and kaempferol, from the flavonoid biosynthesis pathway (Table S1). HPLC analysis indicated that MdUGT83L3 could only catalyze three compounds: cyanidin, quercetin and kaempferol (Figure 2A). In comparison, the specific enzyme activity of MdGT1 to cyanidin is about 3.47 nkat·mg^-1^, which is higher than the specific enzyme activities of quercetin and kaempferol, and has no activity towards others (Table S1). Moreover, the reaction products of MdUGT83L3 in catalyzing the cyanidin, quercetin and kaempferol were analyzed by LC-MS, and the corresponding glycosides with a glucose addition were identified (Figure 2B).

**Figure 2.**
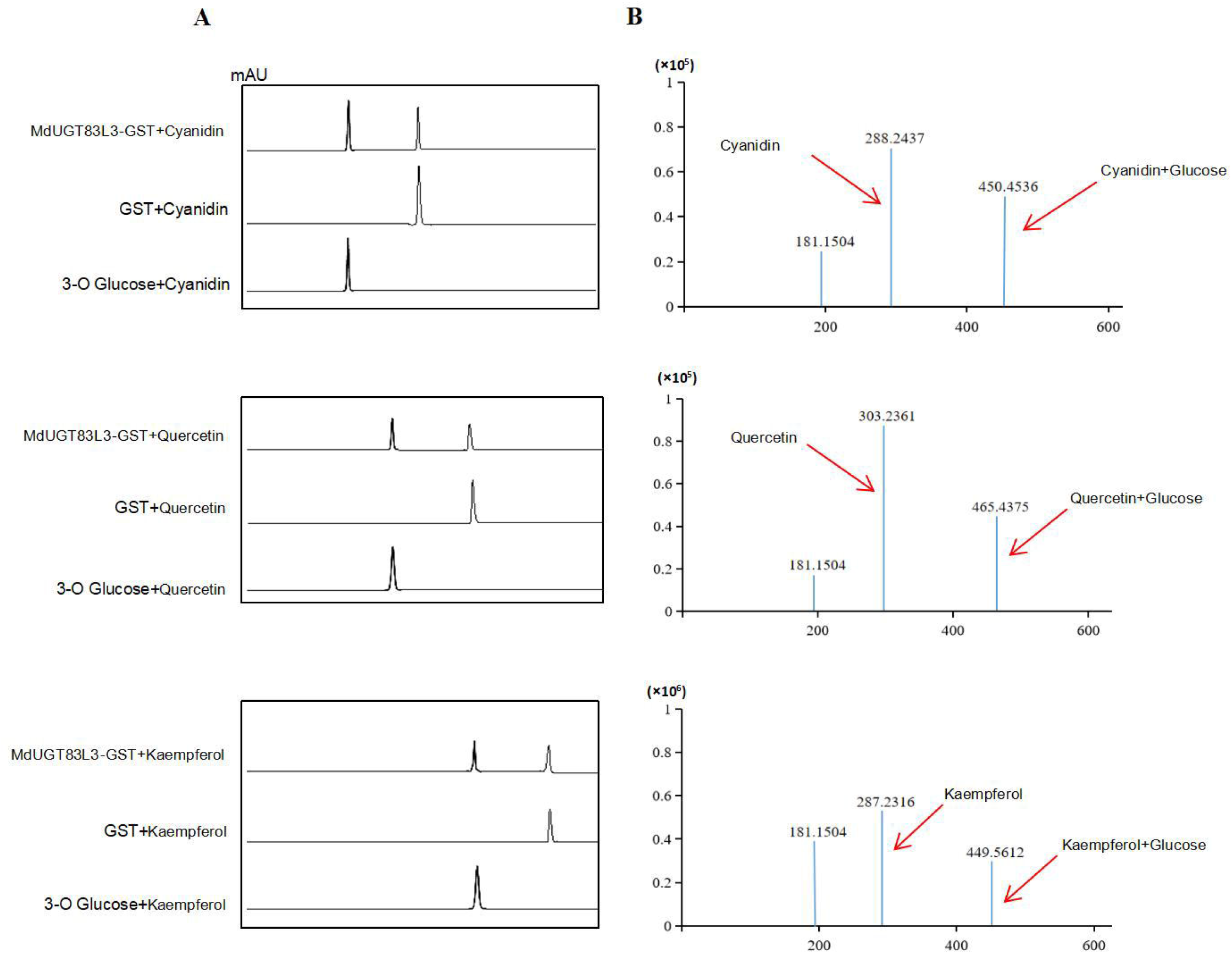
MdUGT83L3 catalyzes the glycosylation of Cyanidin, Quercetin and Kaempferol. **A**. HPLC analysis of the reaction products from cyanidin, quercetin and kaempferol. **B**. LC-MS confirmation of reaction products from cyanidin, quercetin and kaempferol.

### Overexpression of *MdUGT83L3* in apple callus enhances anthocyanin accumulation and stress tolerance

Besides abiotic stresses, it is also observed that *MdUGT83L3* is responsive to cyanidin, quercetin and kaempferol, and is more induced by cyanidin than the other two substrates (Figure 3A). Based on the above findings, we further investigated the function of *MdUGT83L3* in anthocyanin accumulation and stress tolerance, The gene was overexpressed in apple fruit callus, and three transgenic lines OE1, OE2, and OE3 with high expressional abundances (above 20 folds) were selected and used for subsequent experiments (Figure 3B). Here, we focused on examining *MdUGT83L3* roles in resisting salt and cold stress. As shown in Figure 4C, under normal condition, the growing of the callus showed no difference between the OE lines and the lines transformed with empty vector control (VC). However, upon exposure to NaCl, OE1, OE2, and OE3 calluses performed better growing and showed more fresh weight than that of the VC line (Figure 3C, D). The same results were also observed when exposed to cold (4°C).

**Figure 3.**
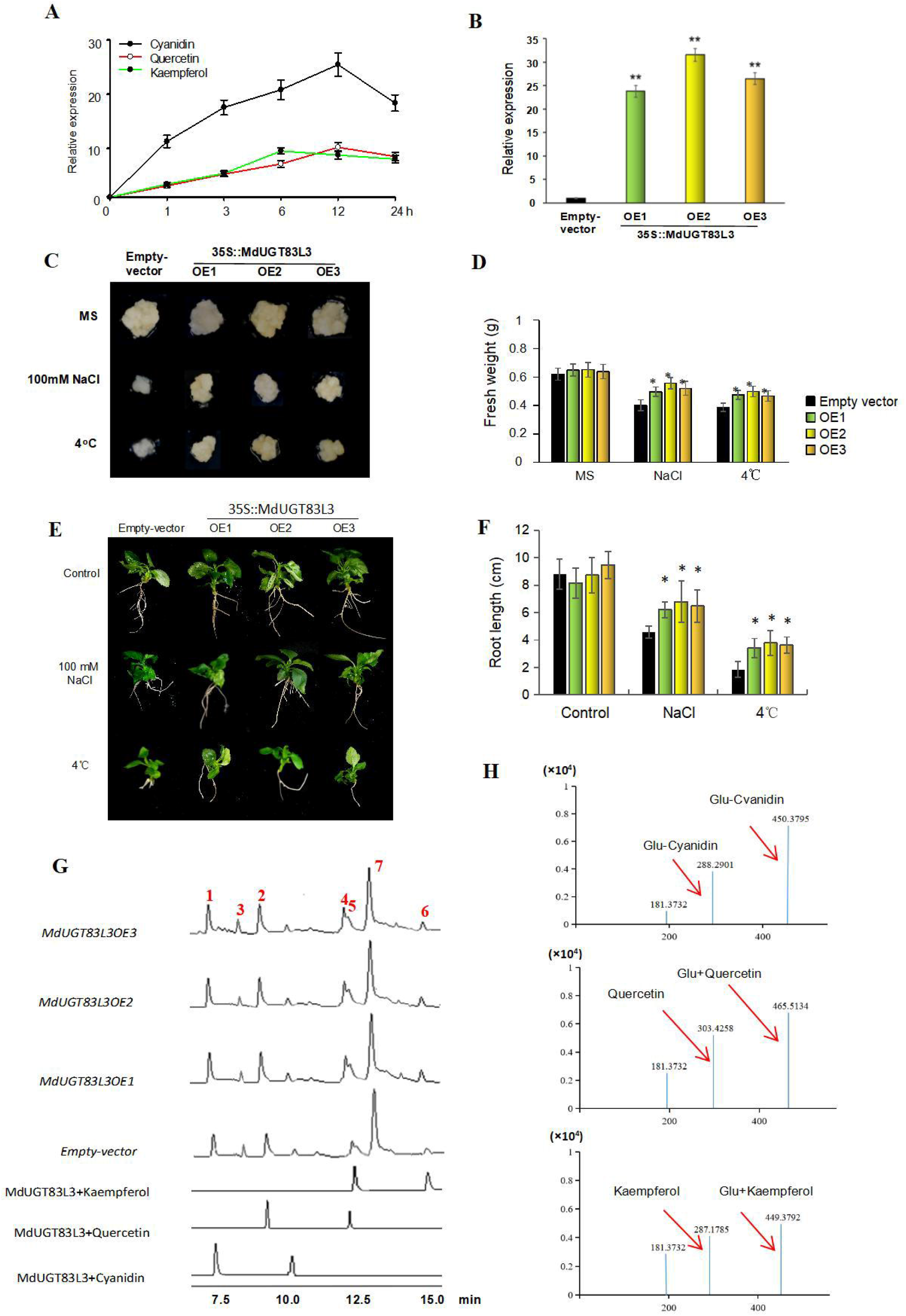
Overexpression of MdUGT83L3 in the callus enhances stress tolerance. **A**. MdUGT83L3 is responsive to cyanidin, quercetin and kaempferol treatment. **B**. MdUGT83L3 expressional levels in the three overexpression lines. **C**. MdUGT83L3 overexpression enhances stress tolerance in callus. **D**. The MdUGT83L3OE lines increased more fresh weight upon exposure to stress conditions. **E**. The regenerated apple plants with uniform growth were subjected to NaCl and cold treatment on the medium for two weeks and then the phenotypes were observed. **F**. The root lengths of the regenerated apple plants were measured. **G**. HPLC analysis of endogenous levels of cyanidin-glucoside, quercetin-glucoside and kaempferol-glucoside in MdUGT83L3OE and control callus (transformed with empty vector). 1:Glucose-cyanidin; 2:Cyanidin; 3:Glucose-Quercetin; 4: Quercetin; 5: Glucose-kaempferol; 6:Kaempferol; 7:IBA (internal standard). **H**. LC-MS confirmation of endogenous cyanidin-glucoside, quercetin-glucoside and kaempferol-glucoside in the callus.

**Figure 4.**
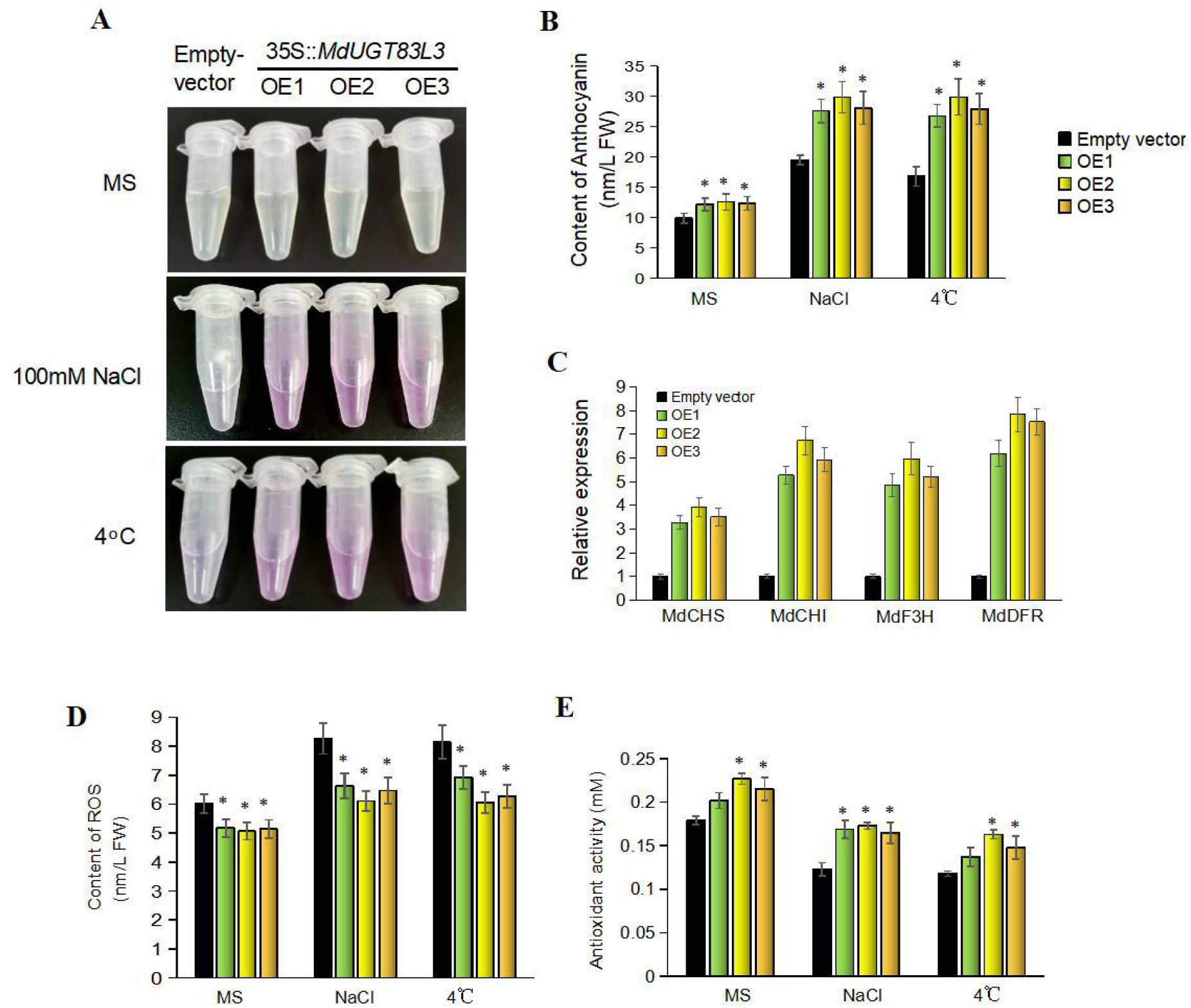
MdUGT83L3 overexpression enhanced anthocyanin accumulation and ROS scavenging in response to NaCl and cold. **A**. Anthocyanin visualized by the purple coloration from NaCl /cold-treated callus. **B**. Anthocyanin contents evaluated by spectrophotometry at a wavelength of 350 or 530 nm. **C**. Anthocyanin biosynthesis pathway genes are upregulated upon MdUGT83L3 overexpression. **D**. ROS content declined in MdUGT83L3 overexpression lines. **E**. Antioxidant activity is enhanced in MdUGT83L3 overexpression lines.

Furthermore, the regenerated seedlings overexpressing MdUGT83L3 were also obtained and subjected to NaCl and Cold treatment. It is found that under NaCl and cold condition, the root length was more increased upon MdUGT83L3 overexpression (Figure 7). These findings also reinforced that MdUGT83L3 can enhance salt and cold stress tolerance in apple.

To investigate the endogenous product catalyzed by MdUGT83L3, the total flavonol were extracted from the *MdUGT83L3OE* callus lines, and analyzed by HPLC and LC-MS. It is observed that the peaks representing Cyanidin glycoside, keampeferol glycoside and quercetin glycoside were obviously elevated in MdUGT83L3OE lines, compared with that of WT (Figure 3G,H).

### Overexpression of MdUGT83L3 enhanced anthocyanin accumulation and removal of ROS

To determine the anthocyanin levels in OE lines, the anthocyanin was extracted and examined by HPLC. The results showed that the anthocyanin was overaccumulated in the overexpression lines and accumulated more under stress conditions than that under control conditions (Figure 4A,B). Consistent with the above results, the expression of the genes involved in anthocyanin biosynthesis including *MdCHS, MdCHI, MdF3H* and *MdDFR* were detected and showed higher expression in the OE calluses (Figure 4C). Flavonoids are natural antioxidants in plants and can remove reactive oxygen species (ROS). Under normal conditions, the ROS level in the callus is low, while after exposed to NaCl and 4°C, the ROS levels increased, which is more in VC lines (Figure 4D). We also examined the antioxidant capacity in the calluses and found that the OE lines exhibited higher antioxidant activity (Figure 4E). These results show that the tolerance of the OE lines to NaCl is due to more ROS scavenging by anthocyanins.

### Overexpression of MdUGT83L3 affected the expression of stress-related genes

To investigate the possible regulating pathways involving MdUGT83L3 in enhancing stress tolerance, we then detected the transcript levels of the stress-related marker genes. The expression of five genes were detected in *MdUGT83L3* overexpression lines, including MdNHX1, MdSOS1, MdCCA1, MdCSP3 and MdCOR47 (Figure S2). Of these, MdNHX1 and MdSOS1 were closely related to salt stress. MdCCA1 and MsCSP3 were involved in cold stress. MdCOR47 was responsive to both salt and cold stress. After dectection, all the selected genes were upregulated upon MdUGT83L3 overexpression in response to salt or cold stresses.

### *MdUGT83L3* is directly regulated by MdMYB88

Next, we want to see how *MdUGT83L3* is regulated in performing its biological roles. Thus, we scanned the 1500bp upstream region of *MdUGT83L3* with the online tool PLACE (http://www.dna.affrc.go.jp/htdocs/PLACE/), and found that its promoter bears a few stress-regulated cis-elements supposed to be bound by MYB transcription factors (Figure 5A). The promoter fragments (P1-P5) containing MYB binding sites were cloned and their interaction with eleven MYB factors were evaluated with yeast one-hybrid assay, including MdMYB111 (LOC103403724), MdMYB46 (LOC103427345), MdMYB74 (LOC103422412), MdMYB306 (LOC103442350), MdMYB20 (LOC103444230), MdMYB58 (LOC103435557), MdMYB88 (LOC103402919), MdMYB44 (LOC103453725), MdMYB59 (LOC103421497), and MdMYB308 (LOC103440814). After these assays, it is found that the only MdMYB88 could bind with the promoter fragment P5, which is the closest element from transcription start site of *MdUGT83L3* (Figure 5B). Additionally, we also performed electrophoretic mobility shift assay (EMSA) to detect the MdMYB88 binding with the P5 fragment and saw the specific shifted band (Figure 5C).

**Figure 5.**
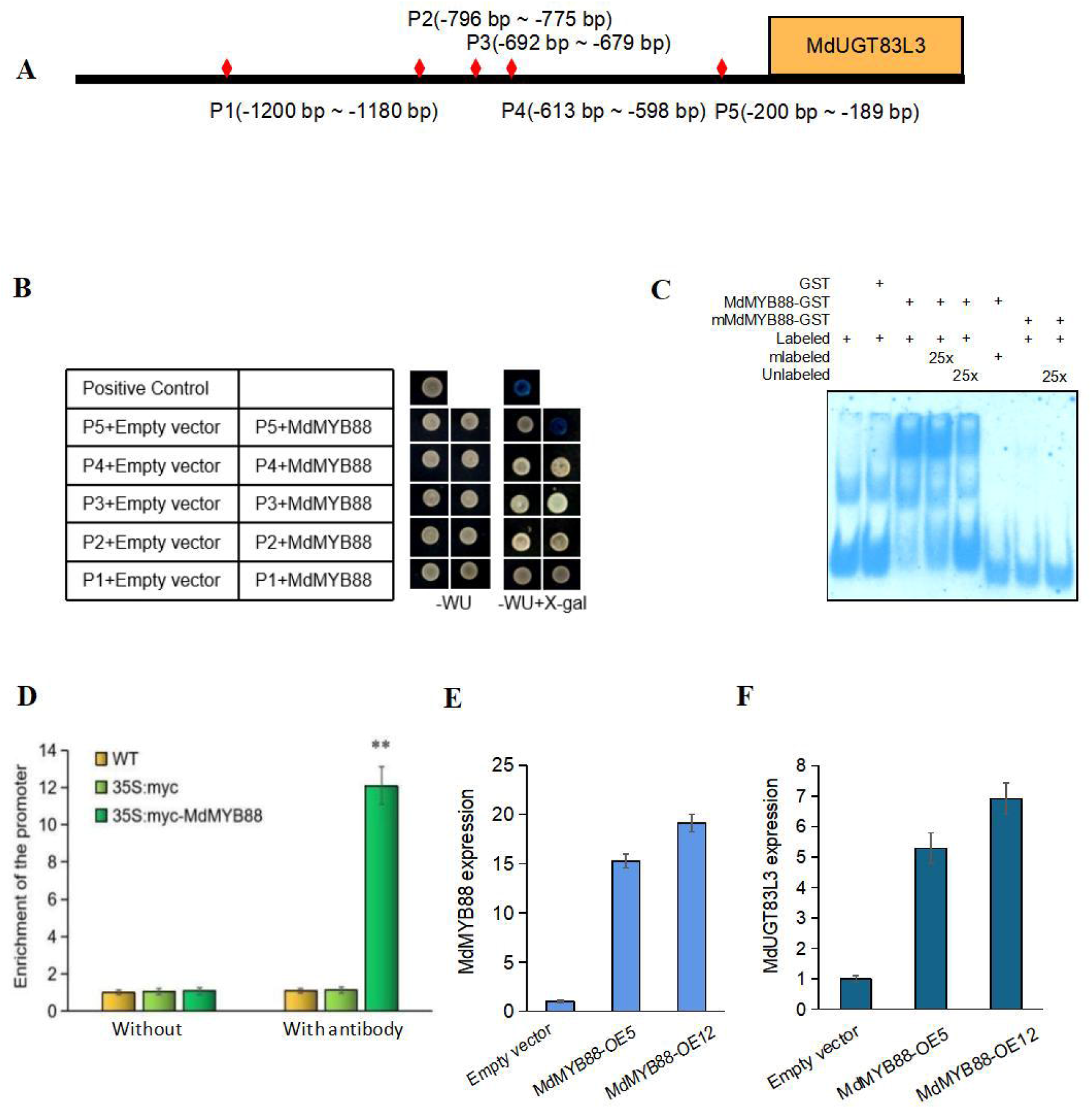
Regulation of MdUGT83L3 expression by MdMYB88 transcription factor. **A**. Promoter fragments (P1-P5) containing the cis-elements in the upstream region of the MdUGT83L3 promoter. **B**. Interaction of MdMYB88 with the promoter fragments evaluated with Yeast-one-hybrid assay. **C**. Interaction of P5 with MdMYB88 assessed by electrophoretic mobility shift assay (EMSA). mlabeled indicates mutated P5 probe. **D**. Enrichment of MdUGT83L3 promoter fragments performed by ChIP quantitative PCR. Before ChIP, all the plants were NaCl-treated for 12 hours, and ChIP was performed using an anti-myc antibody in wild-type (WT) plants, and 35S::myc and 35S::myc-MdMYB88 transgenic plants. Asterisks indicate significant differences relative to the WT (Student’s t-test:*P<0.05; **P<0.01). **E**. Expression levels of MdMYB88 in the apple callus overexpressing MdMYB88. **F**. Upregulation of MdUGT83L3 in MdMYB88 overexpression lines.

Furthermore, to confirm whether CBF1 directly binds to *MdUGT83L3* promoters *in planta*, the chromatin immunoprecipitation (ChIP) assays were then conducted. The myc-tagged MdMYB88 was overexpressed, driven by the CaMV 35S promoter in apple callus. And the transgenic callus were subjected to ChIP assay. After precipitation, we detected by qRT-PCR that the DRE-containing region within the MdUGT83L3 promoters was about 11.8-fold enriched with an anti-myc antibody, respectively (Figure 5D).⍰However, we did not see any obvious enrichment in wild-type callus and the 35S::myc transgenic callus with or without anti-myc (Figure 5D). These findings provide strong evidence that MdUGT83L3 is directly regulated by MdMYB88. Consistent with this notion, the expression of *MdUGT83L3* was significantly elevated in MdMYB88 overexpression calluses (Figure 5E,F).

Based on above findings, we proposed a working model of MdUGT83L3. As shown in Figure 6, in response to salt and cold stresses, MdMYB88 is induced and then activates the expression of *MdUGT83L3* by binding to its promoter, followed by glycosylation of anthocyanin, quercetin and keampeferol, and overaccumulation of these compounds. Anthocyanins have positive roles in scavenging reactive oxygen species, which thereby results in enhanced tolerance to abiotic stresses.

**Figure 6.**
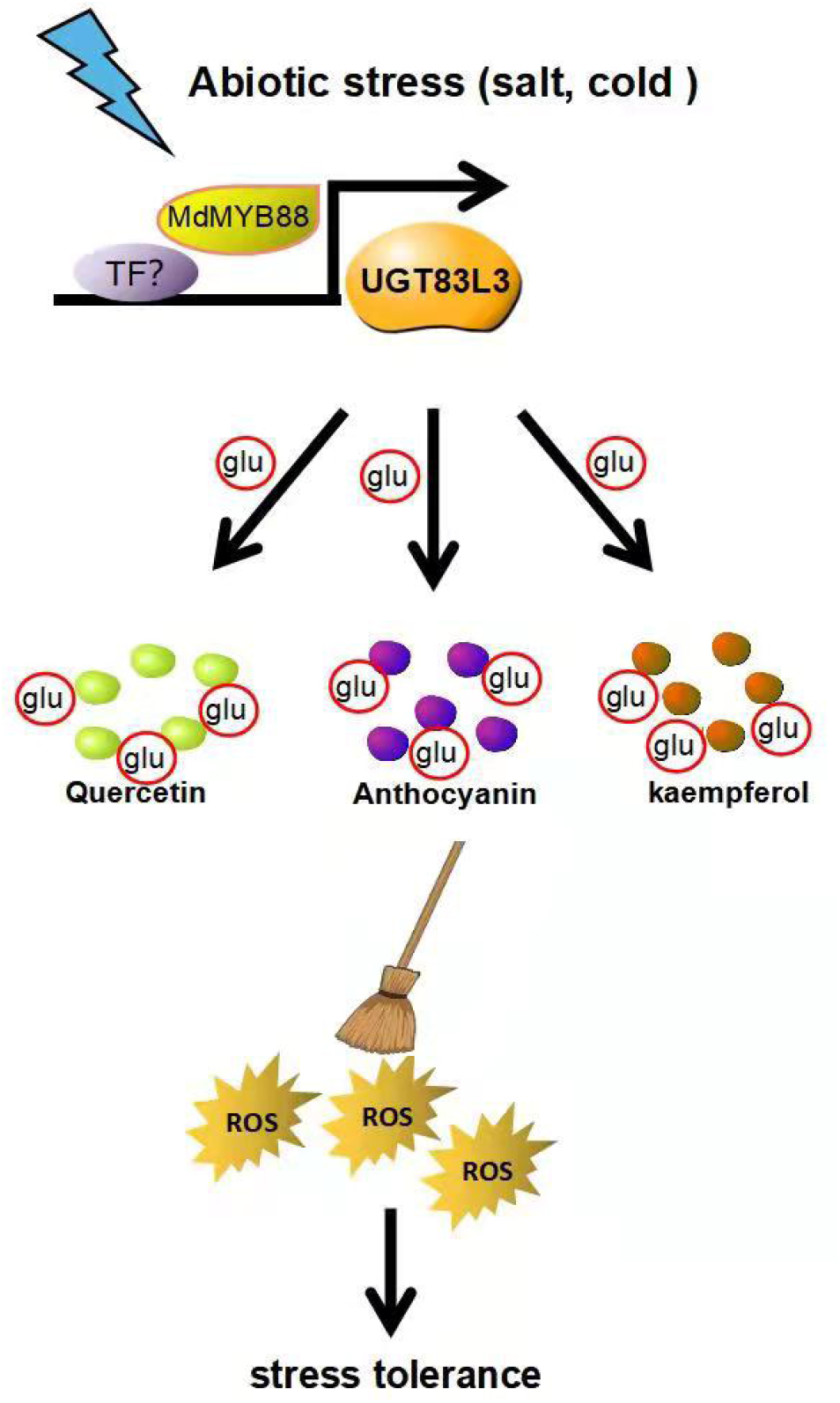
Working model of MdUGT83L3 in response to abiotic stresses. In response to abiotic stresses, MdMYB88 is induced and then activates the expression of MdUGT83L3 by binding to its promoter, followed by glycosylation of anthocyanin, quercetin and keampeferol, and overaccumulation of these compounds. These compounds have positive roles in scavenging reactive oxygen species, which thereby results in enhanced tolerance to abiotic stresses.

## Discussion

Nowadays, the plant UDP-dependent glycosyltransferases (UGTs) have attracted more and more attention duing to their large number and diverse roles. In our study, we identified 229 UGTs in apple through blast against two genome databases (Figure 1A), which is not the same but very close to the previously reported 241 UGTs ^32^. One of the reasons might be due to the update of the genome database. On the other hand, during our screening of the UGTs, it is found that although some sequences have PSPG-box, they also have other domain besides the motif, and the glycosyltransferase activity might just be part of the protein functions. Thus, these protein sequences were removed. In addition, we also removed genes that do not have intact PSPG-box. For some of the sequences that can be alternatively spliced, we just kept one spliced form for these genes. In our study, based on newly identifed UGT groups in the plants ^38,39^, we identified altogether 18 groups (A-R) instead of 16 groups (A-P) in previous study ^32^. Recently, Wilson and Li (2019)^38^ performed detailed analysis to the evolutionary landscape of the plant UGTs. They listed a series of group members from A to R, of which several members were employed in our study to construct the phylogenetic tree. For example, members of group Q and R are clustered with PgUGT95B2 ^39^ and GRMZM2G162783_P01 ^38,40^, respectively. Besides, out of the 229 UGTs, 222 were named, and the rest 7 UGTs are too short to be named in an exact manner (Table 1). It is beleived that the nomenclature will greatly facilitate further study of the UGT functions. Another interesting work in this study is that we discovered many salt/cold stress-induced UGTs after a lot of qRT-PCR analysis (Figure 1C), which provides first information for characterize these genes in salt/cold stress pathways.

During the past two decades, many UGT families have been well-studied in their preferred sugar acceptors, enzyme activities as well as biological roles in diverse plant species. For instance, UGT73 ^41-43^, UGT75 ^37,44^, UGT76 ^41,45^, UGT79 ^10,11^etc. However, functions of many UGTs remain blank, which need to be characterized, such as UGT82, 83, 96, 97, 98, 99 ^38^ etc. In this study, we identified a stress-regulated UGT in apple belonging to group I which is named as MdUGT83L3. It is identified that MdUGT83L3 catalyzes the glycosylation of anthocyanin, quercetin and keampeferol (Figure 2). Also, this gene is significantly induced by salt stress (Figure S1), which is a specific point, because knowledge about salt responsive UGTs is poor. For several flavonoid-modifying UGTs, many of them are stress regulated. For example, UGT79B2 and 79B3 are strongly induced by cold stress. Although they are also responsive to NaCl and mannitol, they might perform a major role in resisting cold stress. Likewise, although MdUGT83L3 also responds to mannitol and cold, it might be more involved in salt stress adaptation. Our study firstly characterizes the UGT83 family, which enriched the knowledge of the plant UGTs.

The MdMYB88 is a well-studied stress-regulated transcription factor which has extensive roles in regulating phenylpropanoids and flavonoids biosynthesis. Many of its target genes have been found, such as COLD SHOCK DOMAIN PROTEIN3 (MdCSP3), CIRCADIAN CLOCK ASSOCIATED1 (MdCCA1) ^27^, MdNCED3 ^30^, MdMYB46 and MdVND6 ^28^. Via directly controlling these target genes, MdMYB88 is involved in regulating cold stress, ABA biosynthesis and lignin biosynthesis under drought stress. These studies also revealed that MdMYB88 is closely related to flavonoid biosynthesis pathway. For instance, Xie et al. (2018)^27^ found that MdMYB88 and MdMYB124 could promote anthocyanin accumulation and H_2_O_2_ detoxification in response to cold. However, the cue from MdMYB88/124 to anthocyanin accumulation is not yet clear. In our study, we identified MdUGT83L3, a new target gene of MdMYB88 (Figure 5). it is found that MdUGT83L3 could significantly increase the anthocyanin accumulation and enhanced ROS scavenging (Figure 4). Based on our results, we hypothesize that the anthocyanin accumulation promoted by MdMYB88/124 overexpression might be via or partly via the regulation to MdUGT83L3. Lately, Geng et al. (2020)^28^ reported that the content of some phenylpropanoids and flavonoids were elevated upon MdMYB88/124 overexpression by targeting MdCM2, which is a crucial enzyme in phenylpropanoids biosynthesis pathway. It is hypothesized that in response to stresses, MdMYB88 is soon upregulated and activate the expression of phenylpropanoid biosynthesis pathway genes, such as MdMYB46, MdVND6 and MdCM2, which will lead to the accumulation of many compounds in the phenylpropanoid pathway such as anthocyanin, quercetin, keampferol etc., which will finally be glycosylated by UGTs to facilitate transport or storage. In our study, we observed that the total contents of quercetin and keampeferol were also increased in MdUGT83L3 overexpression lines (Figure 3G,H). Taken together, it is deduced that MdUGT83L3 might be an important component in the MdMYB88/124 regulated phenylpropanoid/flavoniod biosynthesis pathway. When confronting abiotic stresses, MdMYB88 is significantly induced and accelerated the phenylpropanoid/flavonoid metabolic pathway by activating the critical enzyme genes involved in the pathway. Then numerous secondary metabolites are accumulated, of which most of them will finally be modified by glycosylation. On the other hand, MdMYB88 also activates MdUGT83L3, a flavonoid glycosyltransferase, which transfers anthocyanin, quercetin and keamperferol into their glycosides.

Owing to containing rich flavonoids, it is hypothesized that apple might encode many flavonoid modifying UGTs. Among the several reported apple UGTs till now, such as MdUGT88F subfamily ^33^, MdUGT75B1 and MdUGT71B1 ^35^, all of which were found to catalyze flavonoids. However, only MdUGT88F1 is functionally characterized in apple, which mediate the biosynthesis of phloridzin and participates in the plant development and pathogen resistance in apple trees ^34,46^. Here, we characterized the role of MdUGT83L3 in modifying anthocyanins, quercetin and keampferol both in vitro and in planta, and revealed that it is highly related to salt stress response under the regulation of MdMYB88, which links the gene in an intact pathway regulating flavonoid biosynthesis and stress response. Our study reveals a novel pathway in the adaptation to salt stress in apple, which also provide new information in studying UGT functions not only to horticulture, but also to broad plant biology.

## Materials and methods

### Plant materials and growth conditions

The apple seedlings used were Malus domestica “Gala”. The tissue-cultured seedlings were grown at 24°C under long-day conditions (16 h : 8 h, light : dark), and were sub-cultured every 30 d. Apple callus was cultured in a dark room at 25°C, and sub-cultured every 3 weeks.

### Phylogenetic analysis of the apple UGT genes

Multiple alignments of the maize UGT amino acid sequences were carried out using Clustal X v1.83 program. Phylogenetic analysis was performed using the MEGA-X program with the neighbor-joining algorithm and the bootstrap value set to 1,000 replicates. The neighbor-joining and p-distance methods were used with the pairwise deletion option to deal with gaps in the amino acid sequences.

### In vitro enzymatic reaction and HPLC analysis

The recombinant MdUGT83L3 proteins were expressed in E. coli and purified with Glutathione SepharoseTM 4B (GE Healthcare). The glycosyltransferase activity assay was carried out with the conditions described by Hou et al. (2004) ^41^. The products were analysed by HPLC on a Shimadzu HPLC system (http://www.shimadzu.com) using a 5-lM C18 column (150 9 4.6 mm; Zorbax; Agilent, http://www.agilent.com). A linear gradient with increasing acetonitrile (solvent A) against double-distilled H_2_O (solvent B) at a flow rate of 1 ml/min over 35 min was used. Both solutions contained 0.1% trifluoroacetic acid. HPLC conditions were as follows: 0 min 90% B phase; 20 min 25% B phase; 22 min 90% B phase and 35 min stop. For flavonoid detection, the wavelength was 270 nm. The products were further confirmed by the LC-MS system. The LC-MS analytical instrument was a Surveyor MSQ (USA) single-quadrupole liquid chromatography mass spectrometer. The methods and mobile phases were similar to the HPLC conditions. The mass spectrometer operated in a positive electrospray ionization mode with 50 eV and a probe voltage of 5.0 kV. The dry heater was set to 180°C. The data acquisition and analysis were performed with XCALIBUR 2.0.6.

### Plasmid construction and generation of transgenic apple callus

To construct vector for prokaryotic expression, the coding region of MdUGT83L3 was cloned into the PGEX-2T vector fused with a glutathione-S-transferase (GST) tag. To generate the *35S::MdUGT83L3/35S::MdMYB88* construct, the coding region of MdUGT83L3/MdMYB88 was cloned into the pBI121 binary vector driven by the CaMV 35S promoter. For generation of the transgenic apple callus, before transformation, the required callus was prepared and cultured for 10–15 days after subculture, followed by selection for transformation.

### Determination of anthocyanin accumulation

Anthocyanin contents were evaluated according to Ronchi et al. (1997) ^47^. Briefly, the apple callus after stress treatments were crushed and pooled to obtain three replicates and anthocyanins were extracted in 10 Ml of methanol (containing 1% hydrochloric acid) for 2 hours at 4°Cin dark. And then anthocyanin colors were observed and then determined spectrophotometrically at a wavelength of 350 or 530 nm.

### Electrophoretic mobility shift assay

The possible cis-elements in the upstream region of *MdUGT83L3* and the transcription factors binding with them were predicted and analyzed with bioinformatics methods. After analysis, recombinant MdMYB111 (LOC103403724), MdMYB46 (LOC103427345), MdMYB74 (LOC103422412), MdMYB306 (LOC103442350), MdMYB20 (LOC103444230), MdMYB58 (LOC103435557), MdMYB88 (LOC103402919), MdMYB44 (LOC103453725), MdMYB59 (LOC103421497), and MdMYB308 (LOC103440814) proteins fused with His-tag were expressed in E. coli (BL21/DE3) and purified with Ni-NTA columns. Biotin-labeled DNA probes are listed in Table S2. The specific method was performed in accordance with the EMSA kit instructions (Thermo Scientific, USA, 89818).

### Yeast one-hybrid assay

For yeast one-hybrid assay, the 200bp promoter fragments containing MYB binding sites were cloned into pLacZi2u vector. And the coding region of MdMYB88 was cloned into pJG4-5 vector. The two plasmids were co-transformed into the competent yeast cells and the positive clones were selected on SD (glucose) medium, and the next day, it was replaced with SD (galactose and raffinose) medium. After incubation, the activity of β-galactosidase was measured. The primer sets employed to amplify the *MdUGT83L3* promoter regions for yeast one-hybrid assay were listed in Table S2.

### Chromatin Immunoprecipitation

1.0g samples from 35S::myc-MdMYB88 transgenic apple callus were collected. Chip was performed by PierceTM Agarose ChIP Kit (Thermo Scientific). DNA primers including 5 fragments in *MdUGT83L3* promoter, Each fragment is about 200bp. DNA enrichment was analyzed by qRT-PCR. The primer sets employed to amplify the MdGT1 promoter regions for chromatin immunoprecipitation screening were listed in Table S2.

### Statistical analysis

All experiments were performed as three independent biological replicates, with at least three technical replicates established each time. The significance of differences between each experimental sample and the control sample was tested using Student’s t-test. Differences at *P < 0.05 were considered significant, while differences at **P < 0.01 were considered extremely significant.

### Quantitative real-time-PCR

For quantitative real-time PCR (qRT-PCR), total RNAs were extracted with Trizol reagent (Takara). Reverse transcription reactions were performed with the PrimeScript RT reagent kit with gDNA Eraser (Takara). Real-time PCRs were performed with a Bio-Rad real-time thermal cycling system using SYBR-Green. All reactions were done at least three times. The apple gene expression levels were normalized with the reference gene EF1-α. Primer information for the qRT-PCR assay is included in Supplementary Table S1.

## Acknowledgement

We thank the UGT nomenclature committee for naming the apple UGTs. This project was supported by grants from the National Natural Science Foundation of China (Grant No. 32001982, 31972357, 31801821, 31772254), Shandong Provincial Natural Science Foundation of China (Grant No. ZR2019PC041), National Key Research and Development Program of China (2019YFD1000104, SQ2020YFF0422322). Supported by the Open Project Program (Number) of State Key Laboratory of Crop Biology, SDAU, Taian, Shandong.

## Author contributions

L.S.J., S.Z.Z and Y.J.L designed the experiments. P.L and L.Z performed the experiments. Y.J.L and P.L carried out the analysis. M.H.C contributed to nomenclature of UGTs. J.S., Z.J.S. and L.P.J. provided the plant materials. Y.J.L and P.L. wrote the paper. C.C.Z., H.R.S. and B.K.H supervised the whole study.

## Figure legends

**Figure S1. Expression analysis of MD09G1064900 (underlined) in response to abiotic stresses (A) and in various tissues (B) by quantitative real-time PCR**.

**Figure S2. Expression of stress-regulated genes is upregulated in MdUGT83L3 overexpression lines when subjected to NaCl and Cold treatment**.

**Table S1. Specific activity of MdUGT83L3 to different substrates**.

**Table S2. Primers used in this study**.

