## Supplementary figures and images for "Genome-wide analysis of the apple family 1 glycosyltransferases identified a flavonoid-modifying UGT, MdUGT83L3, which is targeted by MdMYB88 and contributes to stress adaptation"

### Fig S1

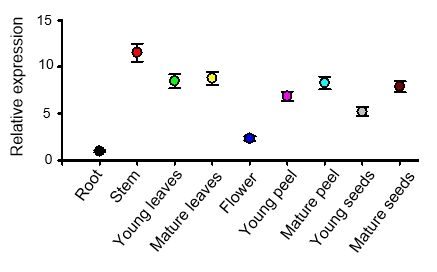

**A**

**B**
