## Supplementary material for "Genome-wide analysis of the apple family 1 glycosyltransferases identified a flavonoid-modifying UGT, MdUGT83L3, which is targeted by MdMYB88 and contributes to stress adaptation": Table S1

***Table 1.* Characteristics of family 1 UGTs from apple.**

| ***Gene ID*** | **Length (aa)** | **Chr.** | **Nomenclature** | **Group** |
| --- | --- | --- | --- | --- |
| MD09G1140700 | 471 | 9 | UGT71A15 | E |
| MD17G1129700 | 474 | 17 | UGT71A63 |  |
| MD09G1141500 | 472 | 9 | UGT71A64 |  |
| MD09G1141100 | 471 | 9 | UGT71A65 |  |
| XP_028963968.1 | 471 | 9 | UGT71A66 |  |
| MD09G1141800 | 488 | 9 | UGT71A67 |  |
| MD09G1141700 | 483 | 9 | UGT71A68 |  |
| MD09G1141300 | 505 | 9 | UGT71A69 |  |
| MD09G1141200 | 414 | 9 | UGT71A70 |  |
| MD17G1129500 | 474 | 17 | UGT71B1 |  |
| MD11G1076200 | 496 | 11 | UGT71K7 |  |
| MD11G1076800 | 481 | 11 | UGT71K8 |  |
| MD11G1076600 | 477 | 11 | UGT71K9 |  |
| XP_008384817.2 | 478 | 11 | UGT71K10 |  |
| MD09G1142000 | 458 | 9 | UGT71W3 |  |
| MD09G1143400 | 359 | 9 | UGT71W4 |  |
| MD05G1246700 | 342 | 5 | UGT71X5 |  |
| MD02G1153200 | 486 | 2 | UGT72A8 | E |
| MD02G1153100 | 457 | 2 | UGT72A9 |  |
| MD02G1153000 | 465 | 2 | UGT72A10 |  |
| MD15G1267300 | 476 | 15 | UGT72A11 |  |
| MD02G1153500 | 479 | 2 | UGT72A12 |  |
| MD14G1122500 | 445 | 14 | UGT72BM3 |  |
| XP_008392435.2 | 491 | 14 | UGT72BM4 |  |
| MD01G1077200 | 473 | 1 | UGT72B46 |  |
| MD07G1146800 | 472 | 7 | UGT72B62 |  |
| RXH73194.1 | 469 | 15 | UGT72BL1 |  |
| MD06G1146400 | 375 | 6 | UGT72BL2 |  |
| MD06G1103300 | 491 | 6 | UGT72BM1 |  |
| MD06G1103400 | 488 | 6 | UGT72BM2 |  |
| MD06G1103600 | 383 | 6 | UGT72BM3 |  |
| MD06G1103500 | 253 | 6 | UGT72BM4 |  |
| MD00G1055100 | 493 | Unknown | UGT73AB13 | D |
| MD05G1085600 | 482 | 5 | UGT73AC7 |  |
| MD00G1046200 | 477 | Unknown | UGT73AR3 |  |
| MD07G1007600 | 478 | 7 | UGT73AR4 |  |
| MD07G1007400 | 474 | 7 | UGT73AR5 |  |
| XP_008375157.2 | 477 | 7 | UGT73AR6 |  |
| MD17G1100000 | 481 | 17 | UGT73B36 |  |
| MD17G1100300 | 482 | 17 | UGT73B37 |  |
| MD05G1086300 | 481 | 5 | UGT73B38 |  |
| MD05G1085800 | 481 | 5 | UGT73B39 |  |
| MD05G1085700 | 276 | 5 | UGT73B40 |  |
| MD12G1104300 | 515 | 12 | UGT73CG21 |  |
| MD12G1104800 | 500 | 12 | UGT73CG22 |  |
| MD05G1085400 | 498 | 5 | UGT73CP3 |  |
| XP_028959262.1 | 472 | 5 | UGT73CQ1 |  |
| XP_028947888.1 | 461 | 14 | UGT74BL2 | L |
| MD15G1146500 | 470 | 15 | UGT74BM6 |  |
| MD02G1003300 | 456 | 2 | UGT74BM7 |  |
| MD15G1146700 | 470 | 15 | UGT74BM8 |  |
| MD16G1086300 | 465 | 16 | UGT74BK10 |  |
| MD15G1357700 | 456 | 15 | UGT74BP1 |  |
| MD10G1111300 | 509 | 10 | UGT74BQ1 |  |
| MD10G1111000 | 460 | 10 | UGT74BQ2 |  |
| MD15G1305500 | 288 | 15 | UGT74BQ3 |  |
| RXH97410.1 | 415 | 5 | UGT74BR1 |  |
| MD04G1019300 | 474 | 4 | UGT75B1 | L |
| MD04G1019400 | 474 | 4 | UGT75L36 |  |
| MD04G1019500 | 481 | 4 | UGT75L37 |  |
| MD04G1019600 | 474 | 4 | UGT75L38 |  |
| MD04G1019700 | 496 | 4 | UGT75L39 |  |
| MD15G1410400 | 462 | 15 | UGT75T5 |  |
| MD07G1208400 | 457 | 7 | UGT76AE1 | H |
| MD01G1148700 | 457 | 1 | UGT76AE2 |  |
| MD01G1143100 | 458 | 1 | UGT76AE3 |  |
| MD07G1209400 | 436 | 7 | UGT76AE4 |  |
| MD01G1144100 | 457 | 1 | UGT76AE5 |  |
| MD07G1210300 | 458 | 7 | UGT76AE6 |  |
| MD01G1142600 | 425 | 1 | UGT76AE7 |  |
| MD03G1245400 | 480 | 3 | UGT76AE8 |  |
| MD07G1208800 | 465 | 7 | UGT76AE9 |  |
| MD01G1143600 | 455 | 1 | UGT76AE10 |  |
| MD01G1142800 | 397 | 1 | UGT76AE11 |  |
| MD01G1143200 | 419 | 1 | UGT76AE12 |  |
| MD00G1075200 | 419 | Unknown | UGT76AF1 |  |
| MD03G1081900 | 451 | 3 | UGT76AF2 |  |
| MD03G1082100 | 451 | 3 | UGT76AF3 |  |
| MD03G1082200 | 463 | 3 | UGT76AF4 |  |
| MD11G1090500 | 452 | 11 | UGT76AF5 |  |
| RXI08878.1 | 593 | 1 | UGT76AG1 |  |

***Table 1. Characteristics of family 1 UGTs from apple (Continued).***

| ***Gene ID*** | ***Length (aa)*** | ***Chr.*** | ***Nomenclature*** | **Group** |
| --- | --- | --- | --- | --- |
| MD16G1266500 | 479 | 16 | UGT78H4 | F |
| MD16G1266400 | 455 | 16 | UGT78S1 |  |
| MD00G1134400 | 248 | Unknown | UGT78S2 |  |
| MD01G1234400 | 483 | 1 | UGT78T1 |  |
| MD07G1306900 | 483 | 7 | UGT78T2 |  |
| MD10G1055200 | 475 | 10 | UGT78U1 |  |
| MD05G1308000 | 459 | 5 | UGT79B60 | A |
| MD10G1287100 | 471 | 10 | UGT79B61 |  |
| MD05G1307900 | 459 | 5 | UGT79B62 |  |
| MD03G1235400 | 460 | 3 | UGT79G9 |  |
| MD11G1256100 | 435 | 11 | UGT79G10 |  |
| MD11G1256300 | 468 | 11 | UGT79G11 |  |
| MD17G1127800 | 473 | 17 | UGT82F1 | N |
| MD09G1137500 | 469 | 9 | UGT82F2 |  |
| MD09G1064900 | 456 | 9 | UGT83L3 | I |
| MD09G1064700 | 471 | 9 | UGT83L4 |  |
| MD17G1058200 | 453 | 17 | UGT83L5 |  |
| MD17G1058400 | 422 | 17 | UGT83L6 |  |
| MD17G1058100 | 455 | 17 | UGT83L7 |  |
| MD09G1065000 | 340 | 9 | UGT83L8 |  |
| MD09G1065400 | 453 | 9 | UGT83K2 |  |
| MD11G1019200 | 486 | 11 | UGT84A78 | L |
| MD10G1115800 | 473 | 10 | UGT84A79 |  |
| MD10G1098500 | 473 | 10 | UGT84A80 |  |
| XP_008366168.2 | 472 | 11 | UGT84A81 |  |
| MD02G1179400 | 477 | 2 | UGT84N2 |  |
| MD17G1125900 | 476 | 17 | UGT85AP6 | G |
| MD17G1126800 | 482 | 17 | UGT85AP7 |  |
| MD09G1142300 | 488 | 9 | UGT85AP8 |  |
| MD17G1125400 | 486 | 17 | UGT85AP9 |  |
| MD17G1126300 | 485 | 17 | UGT85AP10 |  |
| MD17G1125800 | 485 | 17 | UGT85AP11 |  |
| MD09G1142800 | 485 | 9 | UGT85AP12 |  |
| MD09G1142500 | 501 | 9 | UGT85AP13 |  |
| MD09G1136200 | 486 | 9 | UGT85AP14 |  |
| MD00G1121300 | 517 | Unknown | UGT85AP15 |  |
| MD17G1125000 | 488 | 17 | UGT85AP16 |  |
| MD16G1231000 | 492 | 16 | UGT85AP17 |  |
| MD13G1225900 | 479 | 13 | UGT85AP18 |  |
| MD13G1226000 | 484 | 13 | UGT85AP19 |  |
| MD00G1121500 | 393 | Unknown | UGT85AP20 |  |
| MD17G1124900 | 478 | 17 | UGT85AP21 |  |
| MD09G1135900 | 478 | 9 | UGT85AP22 |  |
| MD17G1124400 | 487 | 17 | UGT85AP23 |  |
| MD17G1124800 | 497 | 17 | UGT85AP24 |  |
| MD09G1136300 | 467 | 9 | UGT85AP25 |  |
| MD17G1126900 | 313 | 17 | UGT85AP26 |  |
| MD09G1136100 | 296 | 9 | UGT85AP27 |  |
| MD08G1185500 | 483 | 8 | UGT85A112 |  |
| MD08G1185700 | 484 | 8 | UGT85A113 |  |
| MD15G1371800 | 483 | 15 | UGT85A114 |  |
| XP_008359346.2 | 488 | 8 | UGT85A115 |  |
| MD08G1185000 | 473 | 8 | UGT85A116 |  |
| MD08G1185600 | 479 | 8 | UGT85A117 |  |
| MD01G1033000 | 480 | 1 | UGT85K36 |  |
| MD11G1245700 | 460 | 11 | UGT85K37 |  |
| MD11G1245400 | 493 | 11 | UGT85K38 |  |
| MD01G1029200 | 479 | 1 | UGT85K39 |  |
| MD11G1246000 | 480 | 11 | UGT85K40 |  |
| MD00G1052900 | 483 | Unknown | UGT85K41 |  |
| MD01G1033200 | 569 | 1 | UGT85K42 |  |
| MD09G1284800 | 478 | 9 | UGT85X4 |  |
| MD03G1125900 | 480 | 3 | UGT86A28 | K |
| MD12G1114100 | 477 | 12 | UGT86A29 |  |
| MD12G1114200 | 524 | 12 | UGT86A30 |  |
| MD12G1114000 | 479 | 12 | UGT86A31 |  |
| XP_008349544.2 | 460 | 11 | UGT87Y2 | J |
| MD11G1062600 | 461 | 11 | UGT87Y3 |  |
| MD11G1062700 | 471 | 11 | UGT87AC1 |  |
| MD11G1062300 | 625 | 11 | UGT87H8 |  |
| MD11G1062800 | 465 | 11 | UGT87AC2 |  |
| MD03G1059700 | 312 | 3 | UGT87E9 |  |
| MD11G1062000 | 455 | 11 | UGT87E10 |  |
| MD11G1061800 | 456 | 11 | UGT87E11 |  |
| MD11G1062200 | 571 | 11 | UGT87E12 |  |
| RXI04137.1 | 509 | 3 | UGT87E13 |  |

***Table 1. Characteristics of family 1 UGTs from apple (Continued).***

| ***Gene ID*** | ***Length (aa)*** | ***Chr.*** | ***Nomenclature*** | **Group** |
| --- | --- | --- | --- | --- |
| MD15G1407600 | 483 | 15 | UGT88F1 | E |
| MD15G1407300 | 481 | 15 | UGT88F4 |  |
| MD08G1219100 | 481 | 8 | UGT88F6 |  |
| NP_001315762.1 | 481 | 8 | UGT88F7 |  |
| MD08G1218900 | 480 | 8 | UGT88F8 |  |
| QLF96273.1 | 474 | Unknown | UGT88A32 |  |
| MD01G1210700 | 486 | 1 | UGT88A41 |  |
| MD01G1210600 | 475 | 1 | UGT88A42 |  |
| MD07G1280800 | 475 | 7 | UGT88A43 |  |
| XP_008347244 | 474 | 7 | UGT88A44 |  |
| MD16G1054500 | 473 | 16 | UGT89D22 | B |
| MD16G1054300 | 480 | 16 | UGT89D23 |  |
| MD06G1206400 | 503 | 6 | UGT89K5 |  |
| MD17G1035900 | 480 | 17 | UGT90L1 | C |
| MD17G1036200 | 480 | 17 | UGT90L2 |  |
| MD17G1036700 | 481 | 17 | UGT90L3 |  |
| MD02G1083600 | 455 | 2 | UGT91AJ1 | A |
| MD02G1084000 | 475 | 2 | UGT91AJ2 |  |
| MD15G1211400 | 469 | 15 | UGT91AJ3 |  |
| MD10G1101200 | 469 | 10 | UGT91AJ4 |  |
| MD02G1083100 | 445 | 2 | UGT91AJ5 |  |
| MD02G1083300 | 453 | 2 | UGT91AJ6 |  |
| MD02G1083800 | 469 | 2 | UGT91AJ7 |  |
| RXI06107.1 | 468 | 2 | UGT91AJ8 |  |
| XP_017183030.2 | 454 | 2 | UGT91AJ9 |  |
| MD15G1310300 | 474 | 15 | UGT91A21 |  |
| MD01G1041000 | 280 | 1 | UGT91A22 |  |
| MD03G1019100 | 473 | 3 | UGT91AK1 |  |
| MD00G1112100 | 469 | Unknown | UGT91AK2 |  |
| MD01G1068500 | 508 | 1 | UGT91AK3 |  |
| MD01G1068800 | 473 | 1 | UGT91AK4 |  |
| XP_008346533.2 | 476 | 1 | UGT91AK5 |  |
| MD08G1200700 | 487 | 8 | UGT91C7 |  |
| MD04G1033600 | 513 | 4 | UGT92G11 | M |
| MD04G1029200 | 506 | 4 | UGT92G12 |  |
| MD04G1030200 | 503 | 4 | UGT92G13 |  |
| MD04G1033500 | 507 | 4 | UGT92G14 |  |
| MD04G1029700 | 517 | 4 | UGT92G16 |  |
| MD11G1022200 | 498 | 11 | UGT92R1 |  |
| MD11G1021900 | 489 | 11 | UGT92R2 |  |
| MD08G1134000 | 471 | 8 | UGT93D3 | O |
| MD14G1076500 | 468 | 14 | UGT93D4 |  |
| MD02G1248500 | 464 | 2 | UGT94AF5 | A |
| MD07G1067600 | 460 | 7 | UGT94AF6 |  |
| MD07G1067700 | 461 | 7 | UGT94AF7 |  |
| MD02G1248100 | 457 | 2 | UGT94AF8 |  |
| MD02G1248200 | 454 | 2 | UGT94AF9 |  |
| MD07G1068000 | 457 | 7 | UGT94AF10 |  |
| MD02G1248300 | 419 | 2 | UGT94AF4 |  |
| MD06G1013900 | 583 | 6 | UGT95J1 | Q |
| RXH93190.1 | 489 | 7 | UGT95G2 |  |
| XP_028961281.1 | 475 | 7 | UGT95G3 |  |
| RXH96859.1 | 571 | 6 | UGT95J2 |  |
| MD15G1106200 | 476 | 15 | UGT97C2 | C |
| MD17G1145300 | 484 | 17 | UGT97E1 |  |
| MD16G1064500 | 406 | 16 | UGT708AB1 | R |
| MD17G1055700 | 384 | 17 | UGT708E4 |  |
| RXH69246.1 | 463 | 17 | UGT708H4 |  |
| MD17G1055500 | 247 | 17 | UGT708H5 |  |
| MD04G1140700 | 491 | 4 | UGT709S2 | P |
| MD12G1156200 | 487 | 12 | UGT709S3 |  |
| MD04G1141400 | 421 | 4 | UGT709S4 |  |
| MD04G1140900 | 256 | 4 | UGT709S5 |  |
| MD04G1141300 | 231 | 4 | UGT709S6 |  |
| MD06G1200500 | 469 | 6 | UGT712J1 | I |
| MD15G1322200 | 332 | 15 | NA | G |
| MD01G1143000 | 303 | 1 | NA | H |
| MD07G1007500 | 252 | 7 | NA | D |
| MD11G1090200 | 252 | 11 | NA | H |
| MD01G1142700 | 215 | 1 | NA | H |
| MD10G1216100 | 372 | 10 | NA | E |
| RXH69428.1 | 274 | 17 | NA | C |
