## Supplementary material for "Genome-wide analysis of the apple family 1 glycosyltransferases identified a flavonoid-modifying UGT, MdUGT83L3, which is targeted by MdMYB88 and contributes to stress adaptation": Table S2

| **Table S2 Primers used in this study** | |
| --- | --- |
| **Primer names** | **sequence (5'-3')** |
| **For amplifying full-length cDNA** | |
| cMdUGT83L3-F | ATGAGCAGCCCACATATTTTAGC |
| cMdUGT83L3-R | CTATGATTTCATCCATTCAATA |
| **For protein expression in *E.coli*** | |
| Pe-MdMYB111-F | GATTATGCCTCTCCCGAATTCATGGGAAGAGCTCCGTGCTG |
| Pe-MdMYB111-R | AGAAGTCCAAAGCTTCTCGAGTTAGTGTCCTTCACCAGTAC |
| Pe-MdMYB46-F | GATTATGCCTCTCCCGAATTCATGAGGAAGCCAGAACCCTC |
| Pe-MdMYB46-R | AGAAGTCCAAAGCTTCTCGAGTCAACTTTGGTAGTCAAGAA |
| Pe-MdMYB74-F | GATTATGCCTCTCCCGAATTCATGGGAAGAGCACCTTGTTG |
| Pe-MdMYB74-R | AGAAGTCCAAAGCTTCTCGAGTCACATGAATTCATTAACATC |
| Pe-RVE8-F | GATTATGCCTCTCCCGAATTCATGCACATTCATTCATTCAG |
| Pe-RVE8-R | AGAAGTCCAAAGCTTCTCGAGCTACTGTTTTCGTAGGACATC |
| Pe-MYB306-F | GATTATGCCTCTCCCGAATTCATGGGGAGGCCTCCTTGCTG |
| Pe-MYB306-R | AGAAGTCCAAAGCTTCTCGAGTCAGAAAAAGTTAGCATTTC |
| Pe-MdMYB20-F: | GATTATGCCTCTCCCGAATTCATGGGAAGGAAACCTTGCTG |
| Pe-MdMYB20-R: | AGAAGTCCAAAGCTTCTCGAGTCACAAGAGCCCATATGCCC |
| Pe-MdMYB58-F | GATTATGCCTCTCCCGAATTCATGGGGGGAAAGGGAAGATC |
| Pe-MdMYB58-F | AGAAGTCCAAAGCTTCTCGAGCTAATTTGAATGGAGGAATTG |
| Pe-MdMYB88-F | GATTATGCCTCTCCCGAATTCATGCCGCAGGAGGAGTCAAAG |
| Pe-MdMYB88-R | AGAAGTCCAAAGCTTCTCGAGTTATAGACTTTGGAGGAGGG |
| Pe-MdMYB44-F | GATTATGCCTCTCCCGAATTCATGGCGGTGATCAGGAAGG |
| Pe-MdMYB44-R | AGAAGTCCAAAGCTTCTCGAGTCACTCAATCCTGCTATACC |
| Pe-MdMYB59-F | GATTATGCCTCTCCCGAATTCATGAAAATGGTGCAAGATC |
| Pe-MdMYB59-R | AGAAGTCCAAAGCTTCTCGAGTCAGCCAGTTAGAGATGC |
| Pe-MdMYB308-F | GATTATGCCTCTCCCGAATTCATGGGAAGGTCTCCTTGCTG |
| Pe-MdMYB308-R | AGAAGTCCAAAGCTTCTCGAGTCATTTCATCTCCAAGCTTC |
| Pe-MdUGT83L3-F | ATGAGCAGCCCACATATTTTAGC |
| Pe-MdUGT83L3-R | CTATGATTTCATCCATTCAATA |
| **For introducing mutation to promoter elements** | |
| DRE-probe-mP5-F | CGATTAATTTAATCAGTATTGGTTAACTCAAATAGTCAGTCAAG |
| DRE-probe-mP5-R | CTTGACTGACTATTTGAGTTAACCAATACTGATTAAATTAATCG |
| **Synthesized probe surrounding proximal DRE element** | |
| DRE-probeP1-F | AATCTTGAATCCGCCAGAATACAAATATCCAATTAGACGGCAACC |
| DRE-probeP1-R | GGTTGCCGTCTAATTGGATATTTGTATTCTGGCGGATTCAAGATT |
| DRE-probeP2-F | GTAATGCTATAGTTAGCCGCTCATACCTAACGACTGTTAATCATT |
| DRE-probeP2-R | AATGATTAACAGTCGTTAGGTATGAGCGGCTAACTATAGCATTAC |
| DRE-probeP3-F | GCGAGCACATATTCGCTCCCTCACAAAAACTACTAAAATTTGGTC |
| DRE-probeP3-R | GACCAAATTTTAGTAGTTTTTGTGAGGGAGCGAATATGTGCTCGC |
| DRE-probeP4-F | GGCAAACAAATGAACAAAGAAACGCTGGTTGGGCTTTTTC |
| DRE-probeP4-R | GAAAAAGCCCAACCAGCGTTTCTTTGTTCATTTGTTTGCC |
| DRE-probeP5-F | CGATTAATTTAATCAGTATTAACCGACTCAAATAGTCAGTCAAG |
| DRE-probeP5-R | CTTGACTGACTATTTGAGTCGGTTAATACTGATTAAATTAATCG |
| **For protein expression in yeast** | |
| PJG4-5-MdMYB111-F | GATTATGCCTCTCCCGAATTCATGGGAAGAGCTCCGTGCTG |
| PJG4-5-MdMYB111-R | AGAAGTCCAAAGCTTCTCGAGTTAGTGTCCTTCACCAGTAC |
| PJG4-5-MdMYB46-F | GATTATGCCTCTCCCGAATTCATGAGGAAGCCAGAACCCTC |
| PJG4-5-MdMYB46-R | AGAAGTCCAAAGCTTCTCGAGTCAACTTTGGTAGTCAAGAA |
| PJG4-5-MdMYB74-F | GATTATGCCTCTCCCGAATTCATGGGAAGAGCACCTTGTTG |
| PJG4-5-MdMYB74-R | AGAAGTCCAAAGCTTCTCGAGTCACATGAATTCATTAACATC |
| PJG4-5-RVE8-F | GATTATGCCTCTCCCGAATTCATGCACATTCATTCATTCAG |
| PJG4-5-RVE8-R | AGAAGTCCAAAGCTTCTCGAGCTACTGTTTTCGTAGGACATC |
| PJG4-5-MYB306-F | GATTATGCCTCTCCCGAATTCATGGGGAGGCCTCCTTGCTG |
| PJG4-5-MYB306-R | AGAAGTCCAAAGCTTCTCGAGTCAGAAAAAGTTAGCATTTC |
| PJG4-5-MdMYB20-F: | GATTATGCCTCTCCCGAATTCATGGGAAGGAAACCTTGCTG |
| PJG4-5-MdMYB20-R: | AGAAGTCCAAAGCTTCTCGAGTCACAAGAGCCCATATGCCC |
| PJG4-5-MdMYB58-F | GATTATGCCTCTCCCGAATTCATGGGGGGAAAGGGAAGATC |
| PJG4-5-MdMYB58-F | AGAAGTCCAAAGCTTCTCGAGCTAATTTGAATGGAGGAATTG |
| PJG4-5-MdMYB88-F | GATTATGCCTCTCCCGAATTCATGCCGCAGGAGGAGTCAAAG |
| PJG4-5-MdMYB88-R | AGAAGTCCAAAGCTTCTCGAGTTATAGACTTTGGAGGAGGG |
| PJG4-5-MdMYB44-F | GATTATGCCTCTCCCGAATTCATGGCGGTGATCAGGAAGG |
| PJG4-5-MdMYB44-R | AGAAGTCCAAAGCTTCTCGAGTCACTCAATCCTGCTATACC |
| PJG4-5-MdMYB59-F | GATTATGCCTCTCCCGAATTCATGAAAATGGTGCAAGATC |
| PJG4-5-MdMYB59-R | AGAAGTCCAAAGCTTCTCGAGTCAGCCAGTTAGAGATGC |
| PJG4-5-MdMYB308-F | GATTATGCCTCTCCCGAATTCATGGGAAGGTCTCCTTGCTG |
| PJG4-5-MdMYB308-R | AGAAGTCCAAAGCTTCTCGAGTCATTTCATCTCCAAGCTTC |
| **MdUGT83L3 promoter fragment for yeast one hybrid** | |
| pLacZi2u-P1MdMYB-F | CGAATACAAATATCCAATTAGG |
| pLacZi2u-P1MdMYB-R | TCGACCTAATTGGATATTTGTATTCGGTAC |
| pLacZi2u-P2MdMYB-F | CGCCGCTCATACCTAACGACTGTG |
| pLacZi2u-P2MdMYB-R | TCGACACAGTCGTTAGGTATGAGCGGCGGTAC |
| pLacZi2u-P3MdMYB-F | CGCACATATTCGCTCCCTCACG |
| pLacZi2u-P3MdMYB-R | TCGACGTGAGGGAGCGAATATGTCGGGTAC |
| pLacZi2u-P4MdMYB-F | CTGGTTGGGCTTTTTCTTTG |
| pLacZi2u-P4MdMYB-R | TCGACAAAGAAAAAGCCCAACCAGGTAC |
| pLacZi2u-P5MdMYB-F | CGTATTAACCGACTCAAATAGG |
| pLacZi2u-P5MdMYB-R | TCGACCTATTTGAGTCGGTTAATACGGTAC |
| **MdUGT83L3 promoter fragment for Chromatin Immunoprecipitation** | |
| Chip-P1MdMYB-F | CGAATACAAATATCC |
| Chip-P1MdMYB-R | TCGACCTAATTGGATAT |
| Chip-P2MdMYB-F | CGCCGCTCATACCTAAC |
| Chip-P2MdMYB-R | TCGACACAGTCGTTAG |
| Chip-P3MdMYB-F | CGCACATATTCGCTCCC |
| Chip-P3MdMYB-R | TCGACGTGAGGGAGCG |
| Chip-P4MdMYB-F | CTGGTTGGGCTTTTTCT |
| Chip-P4MdMYB-R | TCGACAAAGAAAAAGC |
| Chip-P5MdMYB-F | CGTATTAACCGACTC |
| Chip-P5MdMYB-R | TCGACCTATTTGAGTCG |
| Chip-MdMYB88-F | GATTATGCCTCTCCCGAATTCATGCCGCAGGAGGAGTCAAAG |
| Chip-MdMYB88-R | AGAAGTCCAAAGCTTCTCGAGTTATAGACTTTGGAGGAGGG |
| **For real-time PCR** | |
| qMd17G1124900-F | AAAGCTCCTCCACCACGAAG |
| qMd17G1124900-R | ATACCGGAAGTCAGGCAAGC |
| qMd01G1148700-F | GATCGTGTTAAGGACCGGGG |
| qMd01G1148700-R | TCTTCAGGGTCACAATCAGCC |
| qMd17G1126400-F | ATTTTTCCACCTGAGTTTGTTG |
| qMd17G1126400-R | ACTCCTGCAGTAGGCTCTC |
| qMd17G1058100-F | CAAACGCATCGTGAACGCCT |
| qMd17G1058100-R | GTCATTCCTGTCCTCCTTGGG |
| qMdUGT83L3-F | TGCTGATCAGGGTATCGGGT |
| qMdUGT83L3-R | TTTGGTGCCAACTGAATCGC |
| qMd06G1103600-F | CGTGGCACCCTACTTGTTCT |
| qMd06G1103600-R | AATTGGACCGGTTCAGGCAT |
| qMd06G1103400-F | CATTAGACACCCAAGGCCATA |
| qMd06G1103400-R | GGCTAACTCGATGAGGGGAG |
| qMd17G1125900-F | ATCTCTAGCACCCGACTCCC |
| qMd17G1125900-R | ACTGGAGGGGTACTACGGTT |
| qMd09G1141700-F | TTGGAATCACACGCCCTTGA |
| qMd09G1141700-R | CCCACTTCGGTTTCACTGCT |
| qMd06G1103500-F | GATGGGGTCTAGTTGTGCC |
| qMd06G1103500-R | GACCCCCAATGCAACTCTT |
| qMd08G1185700-F | TCAACTCTGCTTCCACCTGT |
| qMd08G1185700-R | CTCAGTCCATAAGTTCGATCCG |
| qMd17G1100000-F | TGGGGAAGCAGAGGAAATGAG |
| qMd17G1100000-R | ACCACGGGATTTCAACTCTTCA |
| qMd06G1200500-F | TTAACCGTTCAGCCCTCACC |
| qMd06G1200500-R | GGCGCCCAAGGAACTATCTT |
| qMd09G1064700-F | TGCCCACCATGAAACCTGAA |
| qMd09G1064700-R | AGGTGAATGTTGCTGGCTCA |
| qMd17G1058200-F | CACAAACGCATCGCCCTGG |
| qMd17G1058200-R | AACTTCCCCGGCATGACTC |
| qMd15G1357700-F | GGTTGCCCTTACTTGAGCCA |
| qMd15G1357700-R | CCATCGCGTTCTCATCCAGT |
| qMd00G1052900-F | AATTGCAGATGGTGTCATGGG |
| qMd00G1052900-R | TGGTGTATCGAGTGTGCCATC |
| qMd17G1126900-F | AGGCCTGACTTGGTAATTGGT |
| qMd17G1126900-R | ACTCCTGCAGACACTTTCCAT |
| qMd09G1141500-F | TGGTGAAGGAGTTGGGACTGT |
| qMd09G1141500-R | TTGCATATCCTTGGCCCGAA |
| qMd01G1210700-F | GCCTTCTAAAAGCCCGAGGT |
| qMd01G1210700-R | GAGGGGAATTGGGCAAGGAA |
| qMd08G1185500-F | TTCGGTTTGAAGCCATCCCA |
| qMd08G1185500-R | TTTGAGGAGGGCACGAAAGG |
| qMd17G1126300-F | CAAAATCCCGGGCTGTAGGT |
| qMd17G1126300-R | GCATATTGTGTTCCGGCCC |
| qMd02G1153200-F | GCAGAGCAAACACCGGAGA |
| qMd02G1153200-R | ACGTGAAGTGTGGATGGGC |
| qMd07G1208800-F | CGTACGGAAAACGCCACCT |
| qMd07G1208800-R | CACAAACCCTCCAGTCACTCT |
| qMd09G1143400-F | CTGTGGTTTTGAACACTTTCGAG |
| qMd09G1143400-R | CTTCTGTCCATAGGCTCGACG |
| qMd08G1185000-F | TGGTGAAGGAGTTGGGACTGT |
| qMd08G1185000-R | GTCAAGCTCCATCACTCGCC |
| qMd09G1065400-F | TTTGTGGGTGGTCAGGGATG |
| qMd09G1065400-R | CCAAGCTACTATCCGCCCAC |
| qMd17G1129500-F | ATGCATTTGAGCTGTTGGAGT |
| qMd17G1129500-R | CCTCTCTCTATTTCTCTGCGCTC |
| qMd09G1136200-F | TTGGCCCTCTTCAGTTGCTT |
| qMd09G1136200-R | GACCACCACACTCCCAAAGT |
| qMd07G1007600-F | CTTCTCGTAGTGTGGCGTTTGT |
| qMd07G1007600-R | CTTTTCCTCGCTCTCCATCGC |
| qMd00G1055100-F | TAGAAGGGGCATGTTCCGGT |
| qMd00G1055100-R | CCGGGACTTCTACACCAACC |
| qMd15G1407300-F | GTTTCAAAGACCTCCGCGAC |
| qMd15G1407300-R | GGCCTGGAGGACGTAGATGA |
| qMd08G1219100-F | AAGGCCTGTGTGTTCCTGAT |
| qMd08G1219100-R | CTCCTCTGCCGCATCTACAAC |
| qMdCHS-F | GGAGACAACTGGAGAAGGACTGGAA |
| qMdCHS-R | CGACATTGATACTGGTGTCTTC |
| qMdCHI-F | GGGATAACCTCGCGGCCAAA |
| qMdCHI-R | GCATCCATGCCGGAAGCTACAA |
| qMdF3H-F | TGGAAGCTTGTGAGGACTGGGGT |
| qMdF3H-R | CTCCTCCGATGGCAAATCAAAGA |
| qMdDFR-F | GATAGGGTTTGAGTTCAAGTA |
| qMdDFR-R | TCTCCTCAGCAGCCTCAGTTTTCT |
| qMdNHX1-F | GCGACAGTCCTGGAACATC |
| qMdNHX1-R | TTATCACTTGCTGCCGGAGG |
| qMdSOS1-F | CGGTTAATCCATCACACACC |
| qMdSOS1-R | TGCTGCCCTGGAGGATTTG |
| qMdCCA1-F | CCACAAGTAGTGGATTTGG |
| qMdCCA1-R | GTTTGCAGTAGAATCAGAG |
| qMdCSP3-F | GTGACAGGACCTAACGGCG |
| qMdCSP3-R | CACCACCTCCACTGCCCTG |
| qMdCOR47-F | GCAGGCGAGACCAAGGATC |
| qMdCOR47-R | CATCACCGGAGATCTTCTCC |
| MdEF-1a-F： | AACTGGTCTGACTACTGA |
| MdEF-1a-R： | TGACAGCAACATTCTTAAC |
